# Dual-localized enzymatic components that constitute the mitochondrial and plastidial fatty acid synthase systems

**DOI:** 10.1101/2020.01.03.894386

**Authors:** Xin Guan, Yozo Okazaki, Rwisdom Zhang, Kazuki Saito, Basil J Nikolau

## Abstract

We report the identification and characterization of genes encoding three enzymes that are shared between the mitochondrial and plastidial-localized Type II fatty acid synthase systems (mtFAS and ptFAS, respectively). Two of these enzymes, β-ketoacyl-ACP reductase (pt/mtKR) and enoyl-ACP reductase (pt/mtER) catalyze two of the reactions that constitute the core, 4-reaction cycle of the FAS system, which iteratively elongate the acyl-chain by 2-carbon atoms per cycle. The third enzyme, malonyl-CoA:ACP transacylase (pt/mtMCAT) catalyzes the reaction that loads the mtFAS system with substrate, by malonylating the phosphopantetheinyl cofactor of acyl carrier protein (ACP). GFP-transgenic experiments determined the dual localization of these enzymes, which were validated by the characterization of mutant alleles, which were transgenically rescued by transgenes that were singularly retargeted to either plastids or mitochondria. The singular retargeting of these proteins to plastids rescued the embryo-lethality associated with disruption of the essential ptFAS system, but these rescued plants display phenotypes typical of the lack of mtFAS function. Specifically, these phenotypes include reduced lipoylation of the H subunit of the glycine decarboxylase complex, the hyperaccumulation of glycine, and reduced growth; all these traits are reversible by growing these plants in an elevated CO_2_ atmosphere, which suppresses mtFAS-associated, photorespiration-dependent chemotypes.

## INTRODUCTION

In plants, *de novo* fatty acid biosynthesis occurs in two distinct subcellular compartments, the plastids and mitochondria (1,2). The two FAS systems utilize an acyl carrier protein (ACP)-dependent, multi-component Type II FAS system, but the two systems use different precursors; acetyl-CoA in the case of plastids, generated by plastidial pyruvate dehydrogenase (ptPDH), and malonyl-CoA in the case of mitochondria, generated by a mitochondrial malonyl-CoA synthetase (3). The plastidial fatty acid synthase (ptFAS) system produces the majority of the plant cell’s fatty acids, and the fatty acids produced by the mitochondrial fatty acid synthase (mtFAS) are also crucial to plant viability (2).

Genetic studies indicate that these two FAS systems are not redundant and have been evolutionarily retained (4). This reflects the alternative fate of the fatty acids generated by each FAS system. The primary role of mtFAS is to synthesize the acyl-precursor for the biosynthesis of lipoic acid (5,6), which is the cofactor essential for the catalytic competence of several key metabolic enzymes, including glycine decarboxylase complex (GDC), mitochondrial pyruvate dehydrogenase (mtPDH), α-ketoglutarate dehydrogenase (KGDH), branched-chain α-ketoacid dehydrogenase (BCKDH) (7), and plastidial PDH (ptPDH) (8,9). In addition, mtFAS appears to be involved in remodeling mitochondrial cardiolipins (10,11), and in detoxifying free malonic acid (3), a competitive inhibitor of succinate dehydrogenase of the TCA cycle (12,13). In contrast, the plastid-originating fatty acids serve as the acyl building blocks for the assembly of the majority of the lipids in plant cells, including membrane lipids, signaling lipids and storage lipids (14).

Typical of Type II FAS systems, the mtFAS system would be expected to be composed of four distinct enzymatic components resembling the Type II FAS that occurs in plant plastids and bacteria (15). These enzymatic components utilize ACP-esterified acyl-substrates to iteratively catalyze a four-reaction cycle, elongating the acyl-chain by 2-carbons per cycle. To date, two of these enzymatic components have been characterized from plant mtFAS, β-ketoacyl-ACP synthase (mtKAS) (16-18) and 3-hydroxyacyl-ACP dehydratase (mtHD) (4). Other supportive components of mtFAS that have been characterized to date, include the mitochondrial phosphopantetheinyl transferase (mtPPT) that activates mtACP by phosphopantetheinylation (19), and malonyl-CoA synthetase (mtMCS) that generates the malonyl-CoA precursor for mtFAS (3). Here in, we report the identification and characterization of three additional enzymatic components. Two of these catalyze two reactions of the mtFAS cycle that have not as yet been identified, namely β-ketoacyl-ACP reductase (KR), and enoyl-ACP reductase (ER), and the third is the mitochondrial malonyl-CoA:ACP transacylase (MCAT), which loads the mtFAS system by malonylating the phosphopantetheinyl cofactor of ACP. Genetic and transgenic expression of fluorescently tagged proteins indicates that these three enzymatic components are shared between mitochondria and plastids.

## RESULTS

### Computational identification of candidate genes encoding mtFAS catalytic components

BLAST analysis of the Arabidopsis genome using the sequences of the *S. cerevisiae* mtFAS components and *E. coli* FAS components as queries, identified putative Arabidopsis ORFs coding for mitochondrial MCAT, KR and ER catalytic components (Supplemental Figure 1). These analyses identified a single candidate each for MCAT (AT2G30200) (20) and KR (AT1G24360) (20), and two potential candidates for ER (AT3G45770 and AT2G05990 (21,22)). The sequences of these candidate proteins are the highest homologs in the Arabidopsis proteome identified in TAIR 10 (www.arabidopsis.org), as indicated by BLAST e-value scores of between 10^−15^ and 10^−68^ and these proteins share between 20% and 45% sequence identities with the query sequences (Supplemental Figure 1).

**Figure 1.**
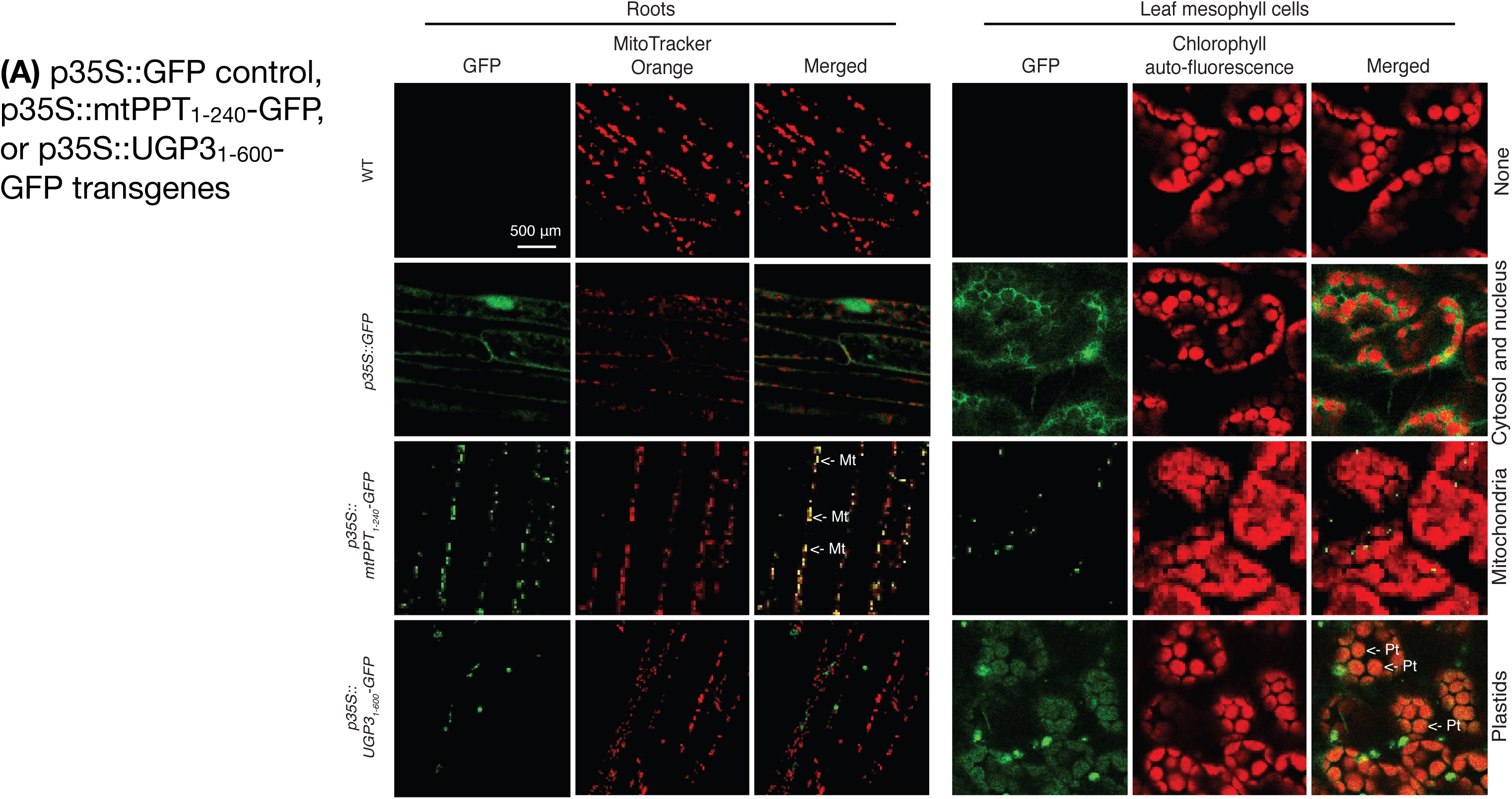
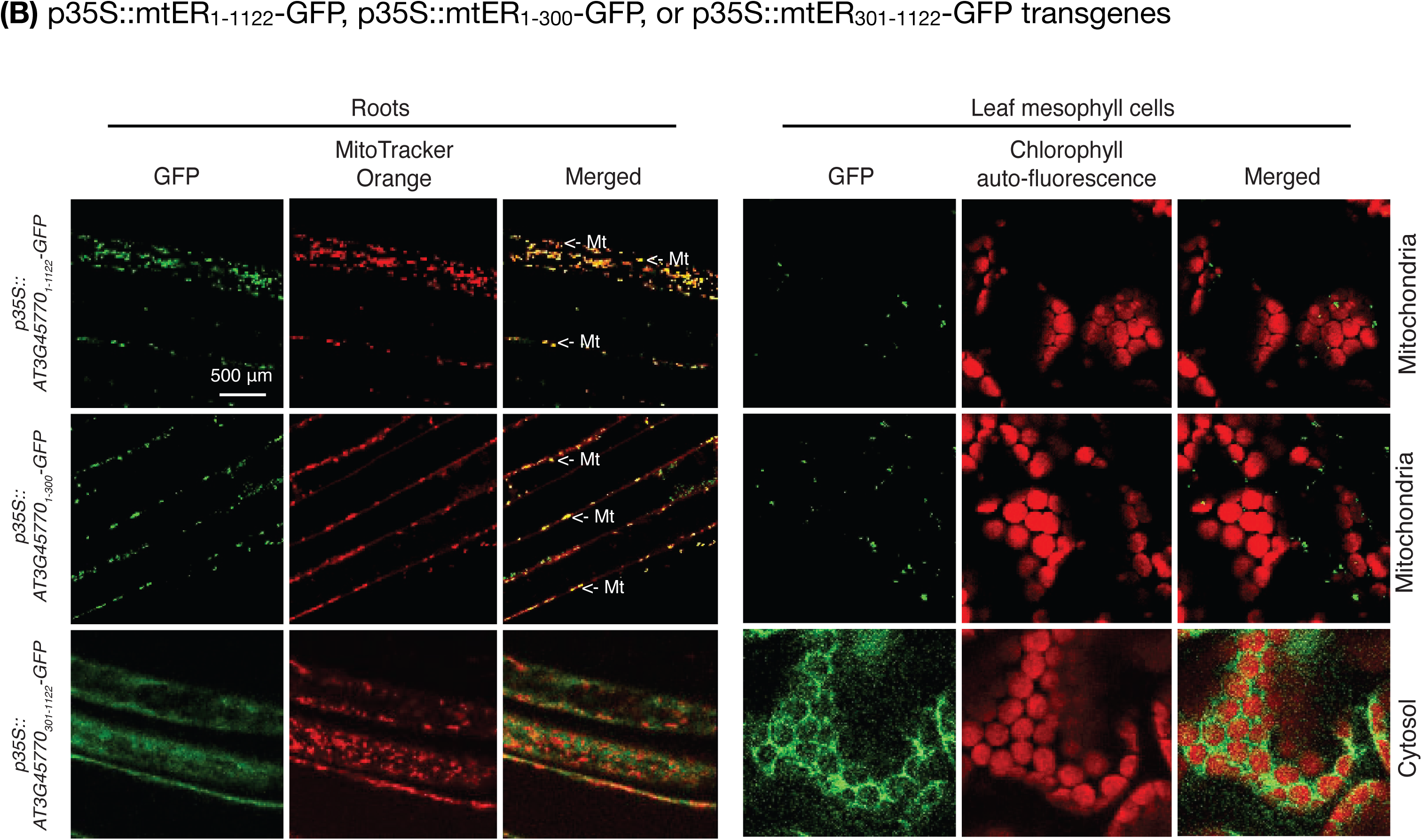
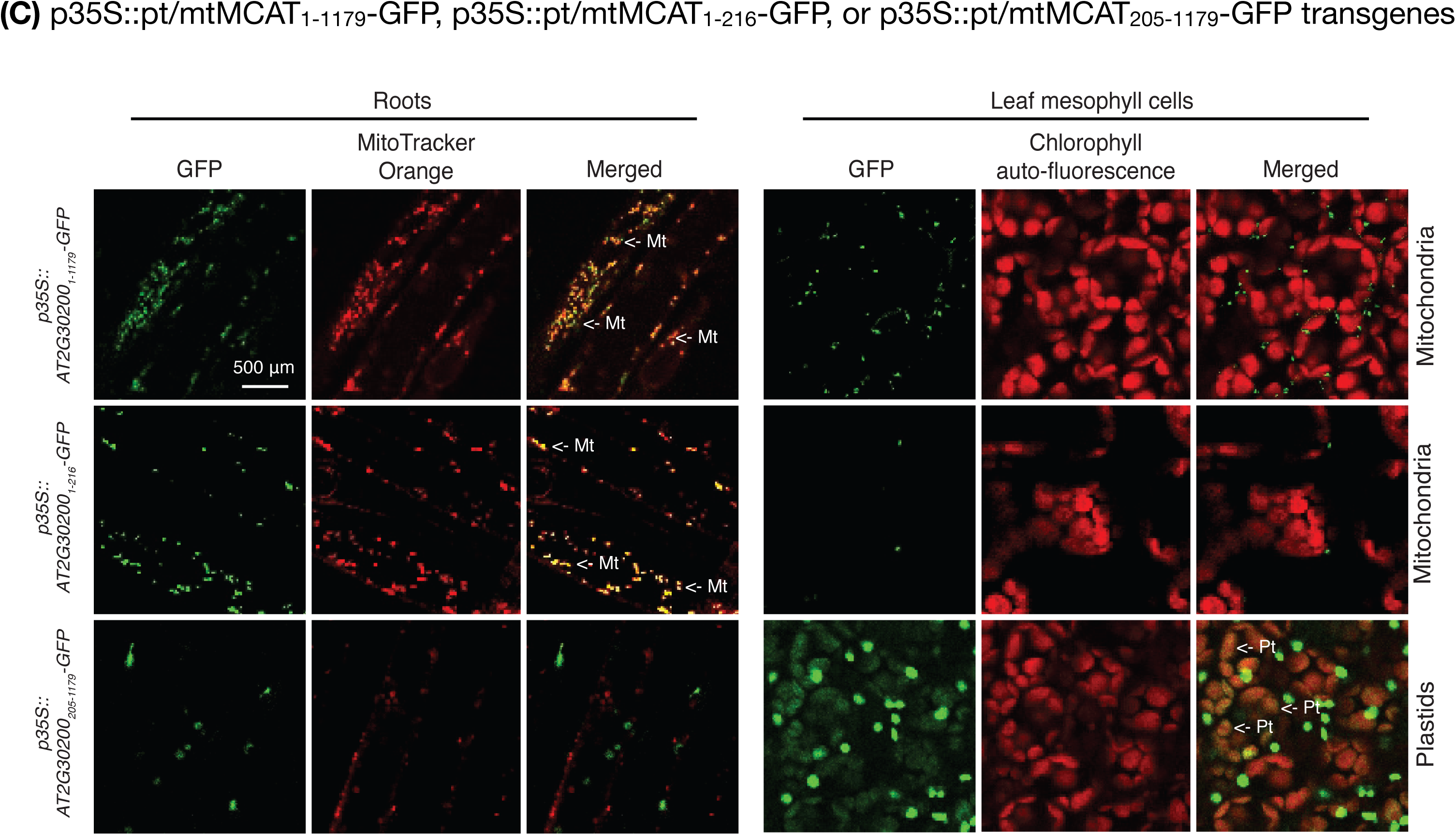
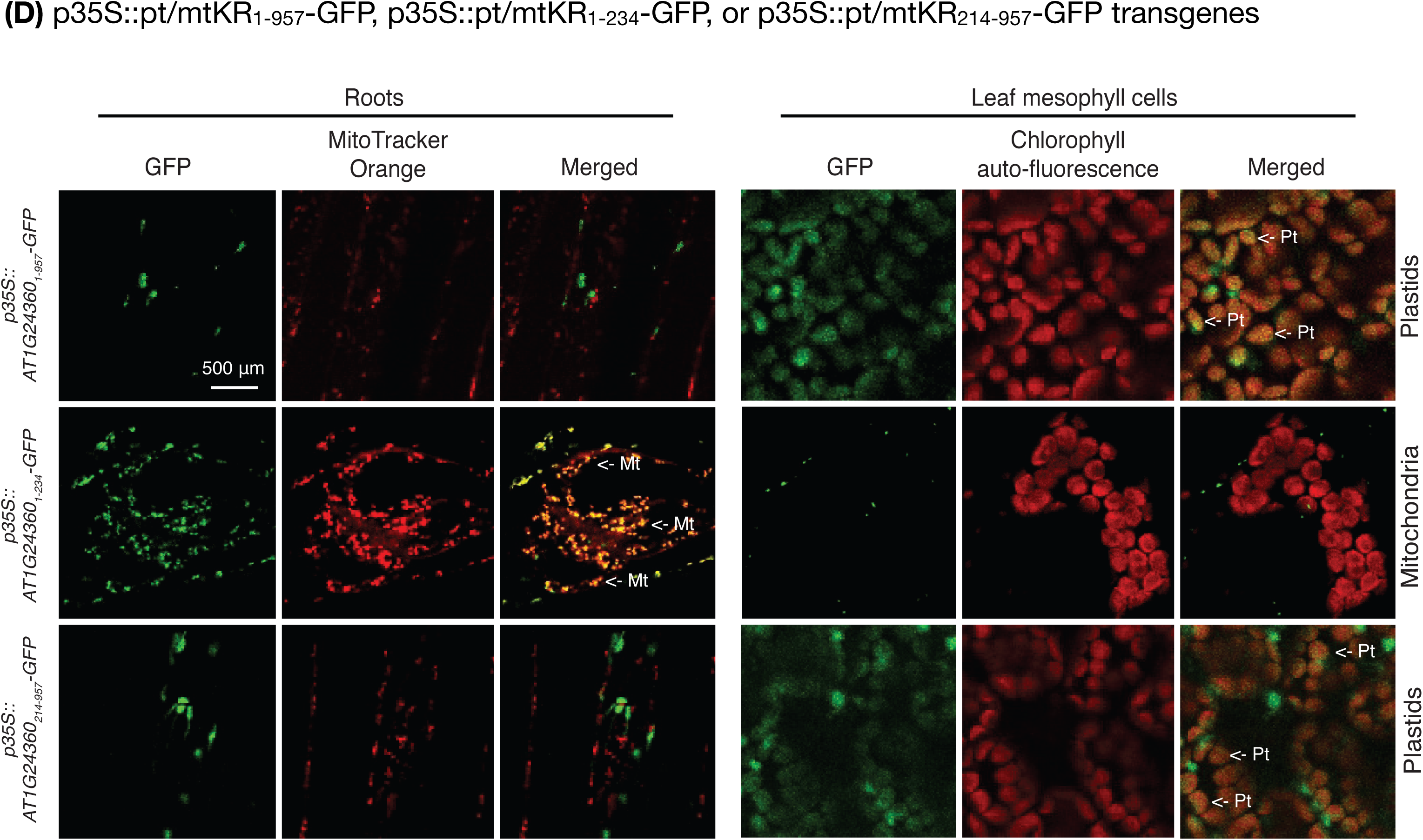
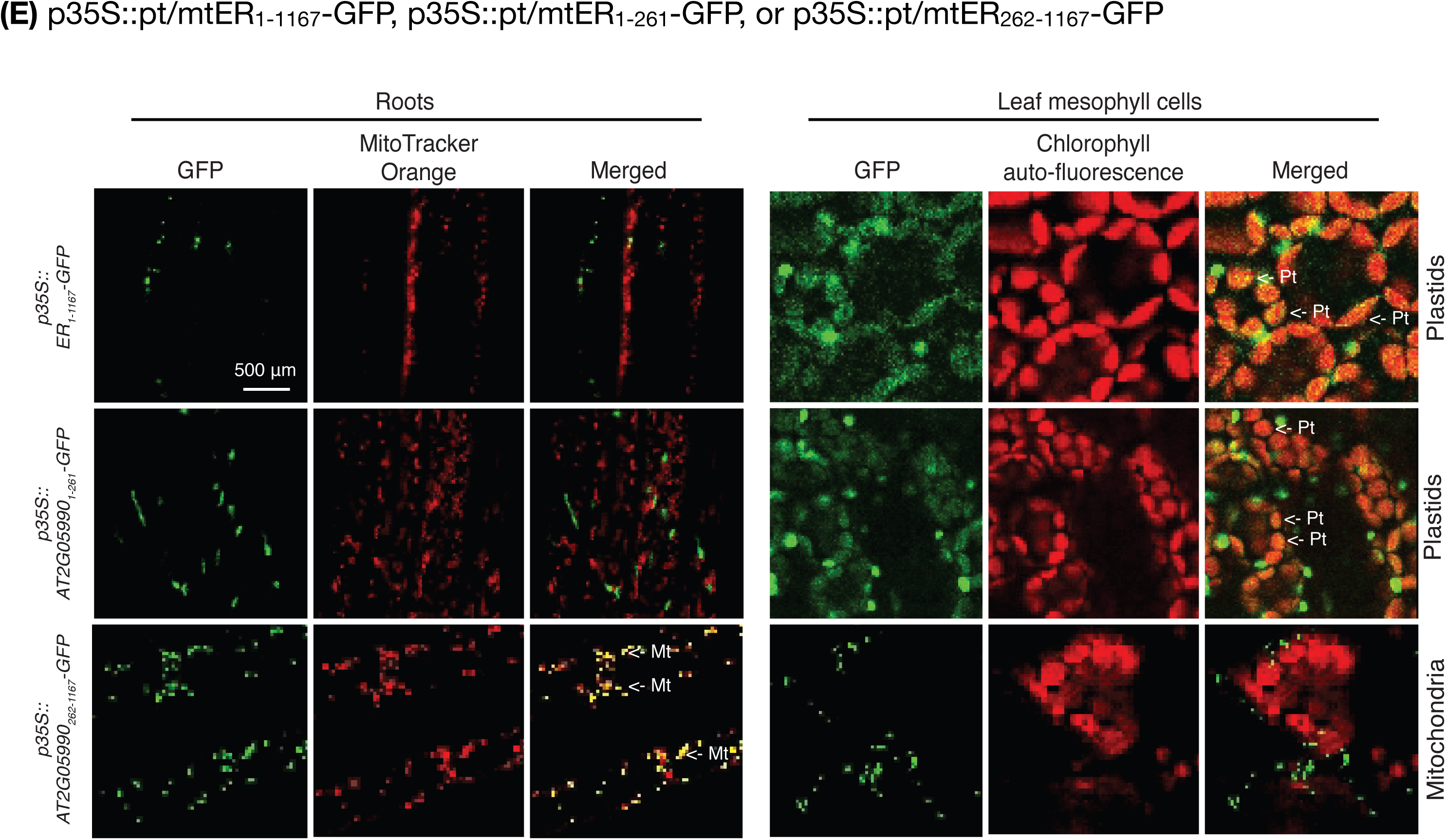
Confocal fluorescence micrographs Confocal fluorescence micrographs of roots and leaf mesophyll cells, imaging the emission of GFP, MitoTracker Orange, chlorophyll auto-fluorescence, and the merged images. Fluorescence micrographs are from non-transgenic wild-type control plants (WT), transgenic plants carrying the *p35S::GFP* control, *p35S::mtPPT_1-240_-GFP*, or *p35S::UGP3_1-600_-GFP* transgenes (**A**); *p35S::mtER_1-1122_-GFP*, *p35S::mtER_1-300_-GFP*, or *p35S::mtER_301-1122_-GFP* transgenes (**B**); *p35S::pt/mtMCAT_1-1179_-GFP*, *p35S::pt/mtMCAT_1-216_-GFP*, or *p35S::pt/mtMCAT_205-1179_-GFP* transgenes (**C**); *p35S::pt/mtKR_1-957_-GFP*, *p35S::pt/mtKR_1-234_-GFP*, or *p35S::pt/mtKR_214-957_-GFP* transgenes (**D**); *p35S::pt/mtER_1-1167_-GFP*, *p35S::pt/mtER_1-261_-GFP*, or *p35S::pt/ mtER_262-1167_-GFP* (**E**).

### Organelle targeting of candidate mtFAS components

Because prior characterizations of these genes indicated that they may be components of the ptFAS system (20-22), initial characterizations evaluated whether these candidate mtFAS components are mitochondrially located. Transgenic experiments were conducted with GFP-fusion proteins, expressed under the transcriptional regulation of the CaMV 35S promoter (Figure 1). Three types of GFP-fusion transgenes were evaluated for each of the candidate mtFAS component proteins: a) GFP was translationally fused at the C-terminus of each candidate mtFAS component proteins; b) GFP was translationally fused at the C-terminus of the N-terminal segment from each candidate mtFAS component protein; these N-terminal segments were computationally predicted to be an organelle-targeting pre-sequence; and c) GFP was translationally fused at the C-terminus of each candidate mtFAS component protein lacking the organelle-targeting pre-sequence. Individual GFP-fusion transgenes were stably integrated into the Arabidopsis genome, and confocal micrographs of the resulting transgenic roots and leaves visualized the subcellular location of the GFP-fusion proteins.

In these experiments, organelles were identified by a combination of two fluorescence markers: a) MitoTracker Orange for mitochondria and chlorophyll auto-fluorescence for plastids; and b) by fluorescence signals revealed by the expression of control GFP-tagged markers, the *p35S::mtPPT_1-240_-GFP* transgene (Figure 1A) that we previously identified as being targeted to mitochondria (19), and the *p35S::UGP3_1-600_-GFP* transgene that is plastid targeted (23) (Figure 1A). These control markers show distinct patterns that are consistent with mitochondrial or plastid localization, and these are distinct from the GFP-signal observed with the non-targeted GFP control, which is located in the cytosol and nucleus, as previously characterized (24) (Figure 1A).

The fluorescence observed in transgenic plants carrying C-terminal GFP-fusions with each full-length candidate mtFAS component-protein show an assortment of organelle localizations. For example, the full-length AT3G45770-encoded protein (rows 1 of Figure 1B) and the N-terminal pre-sequence of this protein (rows 2 of Figure 1B) direct GFP into mitochondria. In contrast, when the N-terminal pre-sequence is removed from the AT3G45770-encoded protein, the GFP is guided to the cytosol (rows 3 of Figure 1B).

The other 3 candidate mtFAS component-proteins guide GFP location to both mitochondria and plastids, with AT2G05990 guiding expression predominantly to mitochondria, whereas AT2G30200 and AT1G24360 guide GFP predominantly to plastids (Figure 1C to 1E). These phenomena were further dissected by fusing the GFP-marker to the N-terminal, potential organelle-targeting pre-sequence of each protein, or fusing the GFP-marker to the C-terminus of each protein that lacks this potential organelle-targeting pre-sequence. These experiments revealed that both the N-terminal, potential organelle-targeting segments, and the mature segments of these proteins contribute to differential targeting of GFP to these two organelles. Specifically, the N-terminal pre-sequences of AT2G30200 (residues 1 to 72) and AT1G24360 (residues 1 to 78) direct the GFP-fusions to accumulate in mitochondria (row 2 of Figures 1C and 1D), but the mature proteins that lack these N-terminal pre-sequences direct the expression of GFP fusion to plastids. However, when these N-terminal pre-sequences are removed from each protein, the remaining segments direct the accumulation of the fused GFP protein to plastids (row 3 of Figures 1C and 1D). These results demonstrate therefore, that the N-terminal pre-sequences of AT2G30200 and AT1G24360 carry mitochondrial-targeting signals. In the case of the AT2G05990, the N-terminal pre-sequence (residues 1 to 87) directs GFP to plastids (row 2 of Figure 1E), but the segment, which lacks this N-terminal pre-sequence, guides GFP to mitochondria (row 3 of Figure 1E).

In summary therefore, proteins encoded by AT2G30200, AT1G24360 and AT2G05990 appear to encode dual localization signals, to both mitochondria and plastids; one of these signals resides in the N-terminal pre-sequence, and the other in the mature portion of these proteins. Furthermore, the signal that resides in the N-terminal pre-sequence appears to predominate in the case of AT2G30200 and AT2G05990, whereas with AT1G24360 the organelle targeting signal in the mature portion of the protein predominates. In contrast, the AT3G45770-protein is singly targeted to mitochondria via a N-terminal pre-sequence signal peptide.

### Experimental authentication of candidate mtFAS component enzymes

The catalytic functions of the individual putative FAS component proteins were explored via two independent strategies. In the first series of experiments, the catalytic function of the putative mtFAS component proteins were evaluated by expressing each protein in *S. cerevisiae* mutant strains that lack a functional copy of a mtFAS component, and testing for genetic complementation. The expression of each candidate Arabidopsis protein was accurately targeted to yeast mitochondria, by genetically fusing the mitochondrial pre-sequence (MP) of the yeast *COQ3* protein (25) to the N-terminus of the mature Arabidopsis proteins, and their expression in yeast was under the transcriptional control of the constitutive PGK promoter (26).

The yeast mutant strains that lack mtFAS functions cannot utilize glycerol as a sole carbon source, because they are deficient in respiration (27). On media that use glycerol as the sole carbon source, the yeast mutant strains lacking individual mtFAS components (i.e., *mct1*, *oar1* or *etr1*) show no growth, unless they express the mitochondrially targeted Arabidopsis proteins encoded by AT2G30200, AT1G24360, AT2G05990 or AT3G45770 (Figure 2A). In each case these results mirrored the results obtained by similarly expressing the yeast MCT1, OAR, or ETR1 proteins (Figure 2A). In contrast, neither the control empty expression plasmid or the expression plasmid carrying only the *COQ3* mitochondrial targeting element is able to rescue the growth deficiency of these yeast strains in glycerol media (Figure 2A).

**Figure 2.**
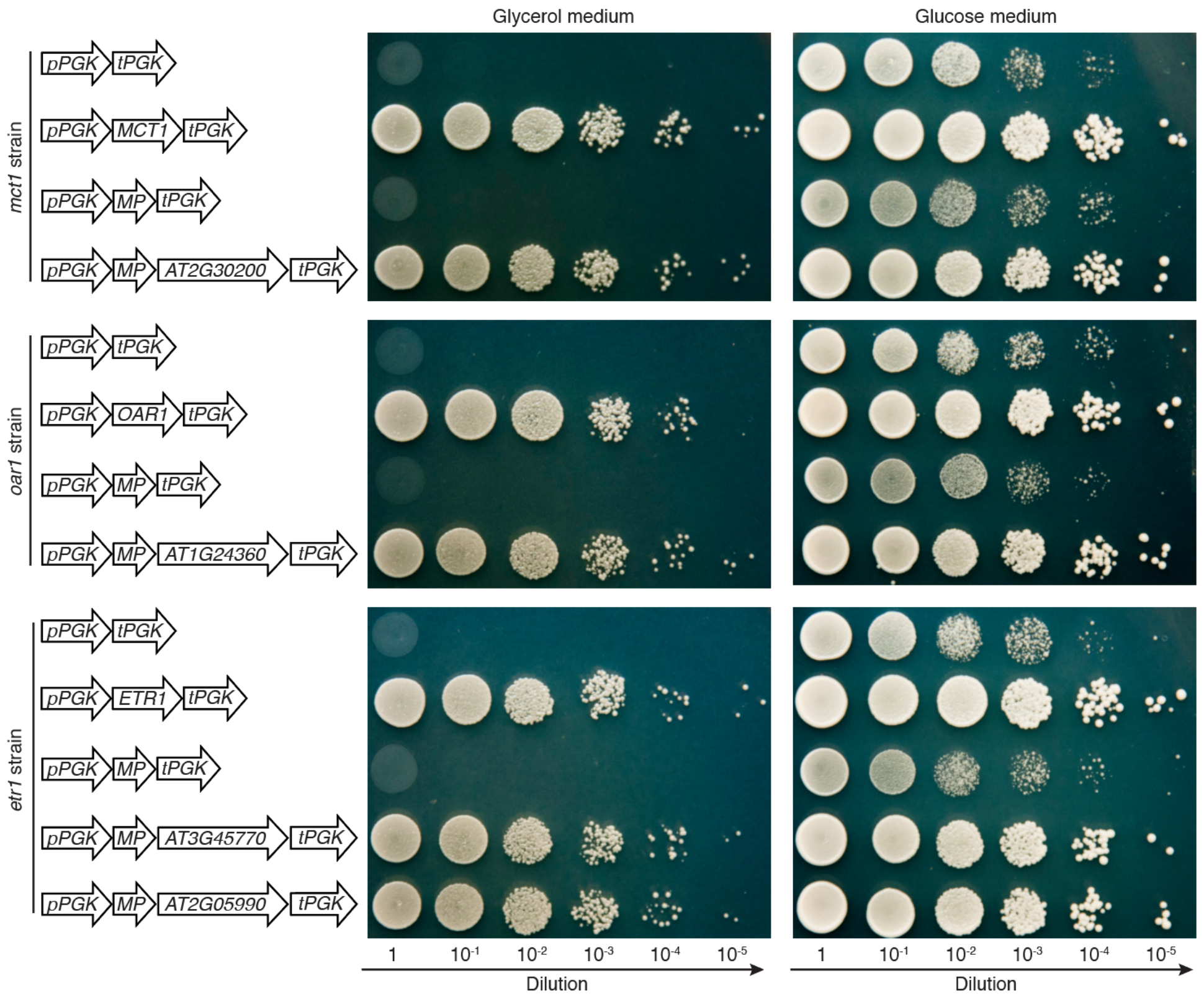
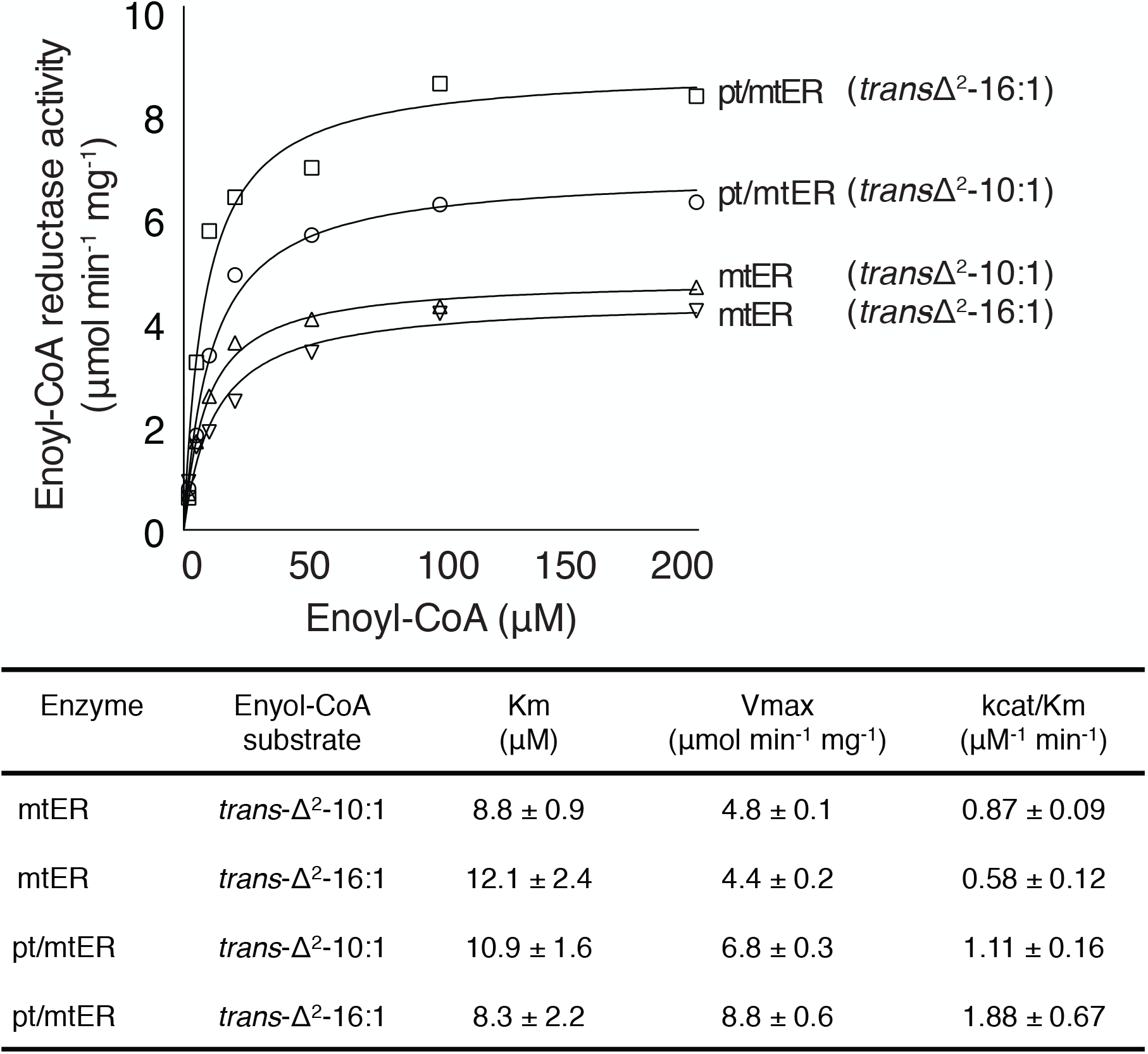
Genetic and biochemical characterization of mtFAS gene candidates **(A)** Genetic complementation of yeast mtFAS mutants (*mct1*, *oar1* and *etr1*) by expression of Arabidopsis mtFAS candidate genes (AT2G30200, AT1G24360, AT3G45770, and AT2G05990); expression of the WT yeast homolog (*MCT1*, *OAR1*, and *ETR1*) served as a positive control. Gene expression was transcriptionally controlled by the phosphoglycerate kinase promoter (*pPGK*) and terminator (*tPGK*). Mitochondrial pre-sequence of yeast COQ3 protein was fused to N-terminus of each protein to ensure the mitochondrial localization. All yeast strains, carrying the indicated expression cassettes were grown on media containing either glycerol or glucose as the sole carbon source, and a dilution series served as the inoculum for each strain. **(B)** *In vitro* characterization of the catalytic capability of purified recombinant Arabidopsis mtER and pt/ mtER enzymes. Substrate concentration dependence of the enoyl reductase activity was assayed with increasing concentrations of either *trans*-Δ^2^-10:1-CoA or *trans*-Δ^2^-16:1-CoA as substrates. The tabulated Michaelis-Menten kinetic parameters were calculated from 3 to 6 replicates for each substrate concentration.

In the case of the enoyl-ACP reductase components of the mtFAS system, recombinant purified proteins encoded by AT2G05990 or AT3G45770 (expressed in *E. coli*) were also evaluated *in vitro* for their ability to catalyze the expected chemical reaction. Because enoyl-ACP reductases are known to be active with both enoyl-ACP (their native substrate) and enoyl-CoA (32), in these experiments each protein was tested for the ability to reduce enoyl-CoA substrates. These assays were conducted with Δ2^*trans*^-10:1-CoA and Δ2^*trans*^ −16:1-CoA, and activity was monitored by the decrease in A340 due to the coupled oxidation of the pyrimidine nucleotides (NADH or NADPH). Both AT2G05990 and AT3G45770 proteins were capable of reducing the enoyl-CoA substrates, and they exhibit comparable K_m_, V_max_ and catalytic efficiency (k_cat_/K_m_) with both tested substrates (Figure 2B). Moreover, these assays establish that AT2G05990 is an NADH-dependent reductase, and its activity with NADPH is undetectable. In contrast, AT3G45770 is an NADPH-dependent reductase, and its activity with NADH is undetectable.

In combination therefore, the GFP-transgenic fluorescence data, the yeast genetic complementation experiments and the biochemical characterizations of purified proteins indicate that three Arabidopsis genes (AT2G30200, AT1G24360 and AT2G05990) encode proteins that are dual targeted to plastids and mitochondria, and they have the ability to catalyze the MCAT, KR and ER reactions, respectively. We therefore label these proteins as pt/mtMCAT (AT2G30200), pt/mtKR (AT1G24360) and pt/mtER (AT2G05990) indicating their dual localizations. In contrast, AT3G45770 encodes a mitochondrially localized ER enzyme, which we label as mtER, indicating its functionality in the sole organelle.

### The *in planta* roles of pt/mtMCAT and pt/mtKR in mtFAS

The role of pt/mtMCAT (AT2G30200) and pt/mtKR (AT1G24360) as enzymatic components of the mtFAS system was further evaluated by characterizing Arabidopsis plants carrying T-DNA-tagged mutant alleles at each locus (details in the method section). As previously described (20), mutant plants homozygous for the T-DNA allele at the AT2G30200 locus are not recoverable as they are embryo lethal. Similarly, mutant plants homozygous for the T-DNA allele at the AT1G24360 also display an embryo lethal phenotype. This embryo-lethality is associated with the fact that these two gene products are components of the ptFAS system (20), which prior genetic studies have established as being essential; these genetic conclusions are exemplified by mutations in other ptFAS components, such as the htACCase (28) and KASI (29).

Therefore, we designed transgenic complementation experiments to further confirm that these dual localized gene products are also components of the mtFAS system. Specifically, plants that are heterozygous mutant at the AT2G30200 locus (pt/mtMCAT) or the AT1G24360 locus (pt/mtKR) were transformed with transgenes that express two versions of the pt/mtMCAT or pt/mtKR proteins, respectively. One version of these transgenes express the ORF that encodes the full-length proteins, and as indicated by the GFP transgenic localization experiments, these full-length proteins contain both an N-terminal, mitochondrial targeting pre-sequence and an internal plastid-targeting signal. These proteins would therefore be expected to be dual-targeted to both plastids and mitochondria. The other version of these proteins express ORFs missing the N-terminal targeting pre-sequence (removing the first 68 and 71 amino acids, respectively, from the full-length proteins) and thereby deleting the mitochondrial-targeting information from each protein. These truncated proteins contain only the internal plastid-targeting signal, and thereby would be expected to express these catalytic functions only in plastids, but not in mitochondria.

Compared to the non-transgenic siblings, which fail to generate homozygous mutant plants (due to missing ptFAS function), all four recovered transgenic lines generated homozygous mutant progeny plants. At 16 day after imbibition (DAI), the transgenic plants, which were expressing the full-length pt/mtMCAT or pt/mtKR proteins were indistinguishable from wild-type plants, indicating that these transgenically expressed proteins complemented the deficiency in both mtFAS and ptFAS function.

In contrast, the transgenic plants expressing the N-terminally truncated pt/mtMCAT or pt/mtKR proteins (i.e., these proteins are predicted to be plastid localized, but not targeted to mitochondria) exhibit reduced size (Figure 3A). Most significantly, when these plants were grown in a 1% CO_2_ atmosphere, where photorespiration deficiency is suppressed, the stunted growth morphology is reversed (Figure 3A). Further biochemical analyses of these plants establish that the lipoylation status of the H protein subunit of glycine decarboxylase (GDC) is reduced to less than 10% of the control levels (Figure 3B). As a consequence of the reduced lipoylation of the H protein, glycine levels are induced by about 100-fold in these plants (Figure 3C). Growing these transgenic mutant plants in the 1% CO_2_ atmosphere restored the accumulation of glycine to near control levels (Figure 3C). Changes in the levels of other amino acids were barely detectable (Supplemental Figure 2).

**Figure 3.**
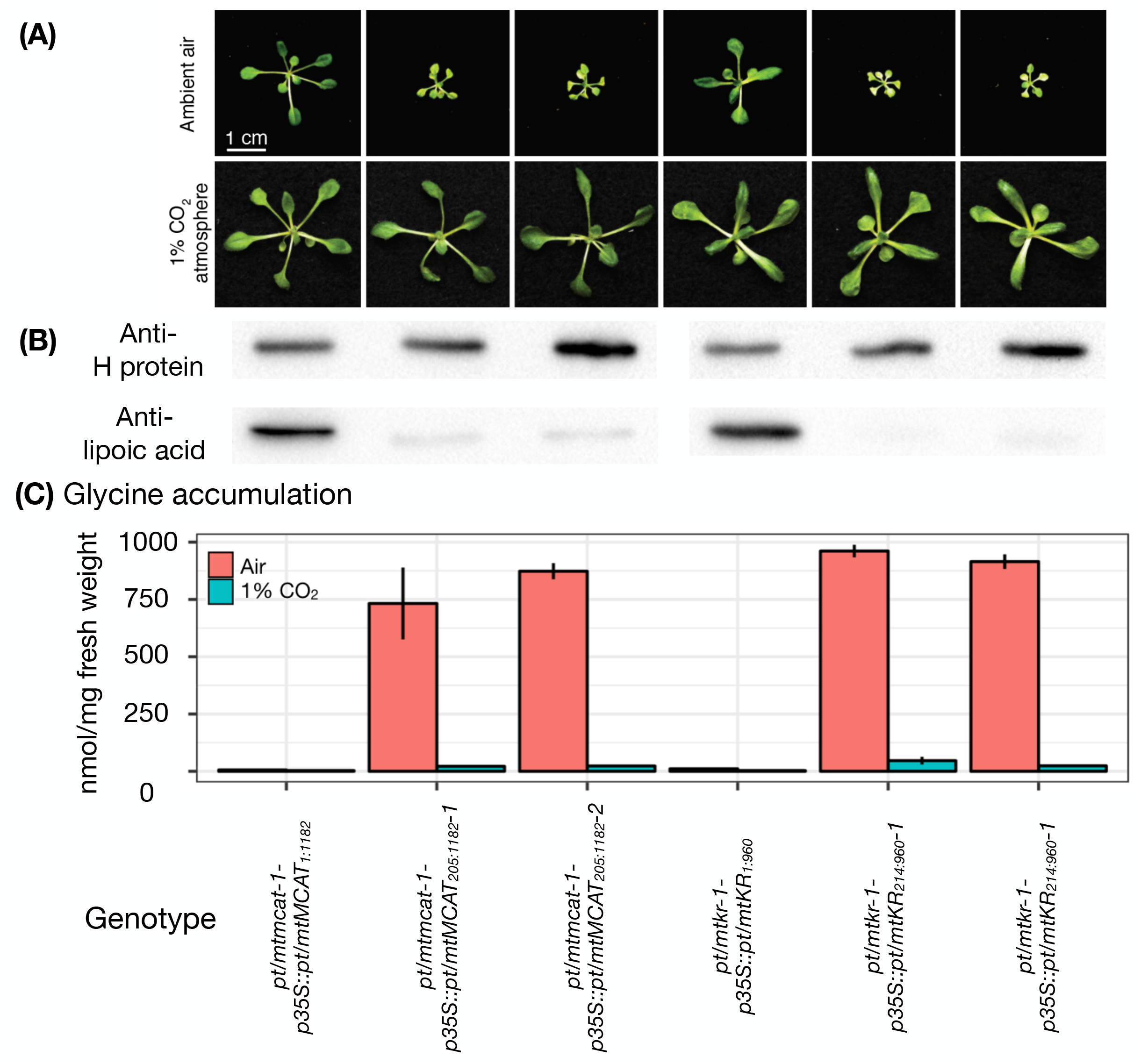
*In vivo* physiological characterization of the *pt/mtMCAT* and *pt/mtKR* genes **(A)** Morphological phenotype of the *pt/mtMCAT* and *pt/mtKR* mutants complemented by full-length or truncated transgenes. Plants were grown in either ambient air or in a 1% CO_2_ atmosphere. **(B)** Western blot analysis of the H subunit of GDC detected with either anti H-protein antibodies, or antilipoic acid antibodies, detecting the lipoylation status of the H-protein. **(C)** Glycine accumulation. Plants of the indicated genotype were grown in either ambient air or in a 1% CO_2_ atmosphere.

These attributes were previously characterized with mutations in other mtFAS components (e.g., mtPPT (19), mtMCS (3), mtKAS (18), and mtHD (4)). These latter mutants exhibit a growth-stunting that is reversible when plants are grown in an elevated CO_2_ atmosphere. Such characteristics have been attributed to the fact that mtFAS generates the lipoic acid cofactor for GDC, and this deficiency blocks photorespiration, leading to the growth deficiency, and hyperaccumulation of glycine, traits that are all reversed when mtFAS mutant plants are grown in an elevated CO_2_ atmosphere that suppresses photorespiration (4). Collectively therefore, these findings demonstrate that the MCAT and KR catalytic functions of the mtFAS and ptFAS systems are genetically encoded by two respective genes that each encode dual localized proteins, pt/mtMCAT and pt/mtKR.

### Two enoyl-ACP reductase isozymes for mtFAS

Two genetic loci appear to encode proteins that catalyze the enoyl-ACP reduction reaction of mtFAS, the AT2G05990 locus that encodes an NADH-dependent reductase and the AT3G45770 locus that encodes an NADPH-dependent reductase. The former protein is dual targeted (i.e., pt/mtER), and the latter is targeted to mitochondria (i.e., mtER). We genetically dissected the roles of these two enoyl-ACP reductase genes by characterizing three mutant lines: 1) two T-DNA-tagged mutant alleles at the mtER locus (*mter-1* and *mter-2*), both of which eliminate the expression of mtER; 2) an RNAi knockdown lines of the *pt/mtER* locus (*pt/mter-rnai-1* and *pt/mter-rnai-2*), which reduce the expression of pt/mtER to ~2% of wild-type levels; and 3) double mutant lines (i.e., *mter-1*-*pt/mter-rnai-1* and *mter-1*-*pt/mter-rnai-2*), which eliminates the expression of mtER and reducing the expression of pt/mtER to ~2% of wild-type levels.

As indicated by the exemplary data gathered from plants at 16 DAI, plants homozygous for the mutant *mtER* alleles are morphologically and metabolically (i.e., amino acids, fatty acids, and lipids) indistinguishable from wild-type plants (Supplemental table 1). In contrast, the aerial organs of the *pt/mter-rnai* mutant and *mter-1-pt/mter-rnai* double mutant lines are considerably reduced in size (Figure 4A). This growth phenotype was equally expressed independent of whether these plants are grown in ambient air or a 1% CO_2_ atmosphere (Figure 4A); the expectation being that in the latter conditions any potential mtFAS associated photorespiration phenotype would be suppressed (18). Therefore, these observations suggest that these mutations do not affect mtFAS, and they are consistent with prior characterization of an ethyl methanesulfonate-generated *pt/mter* mutant (21,22) that identified this gene product as a component of the ptFAS system.

**Figure 4.**
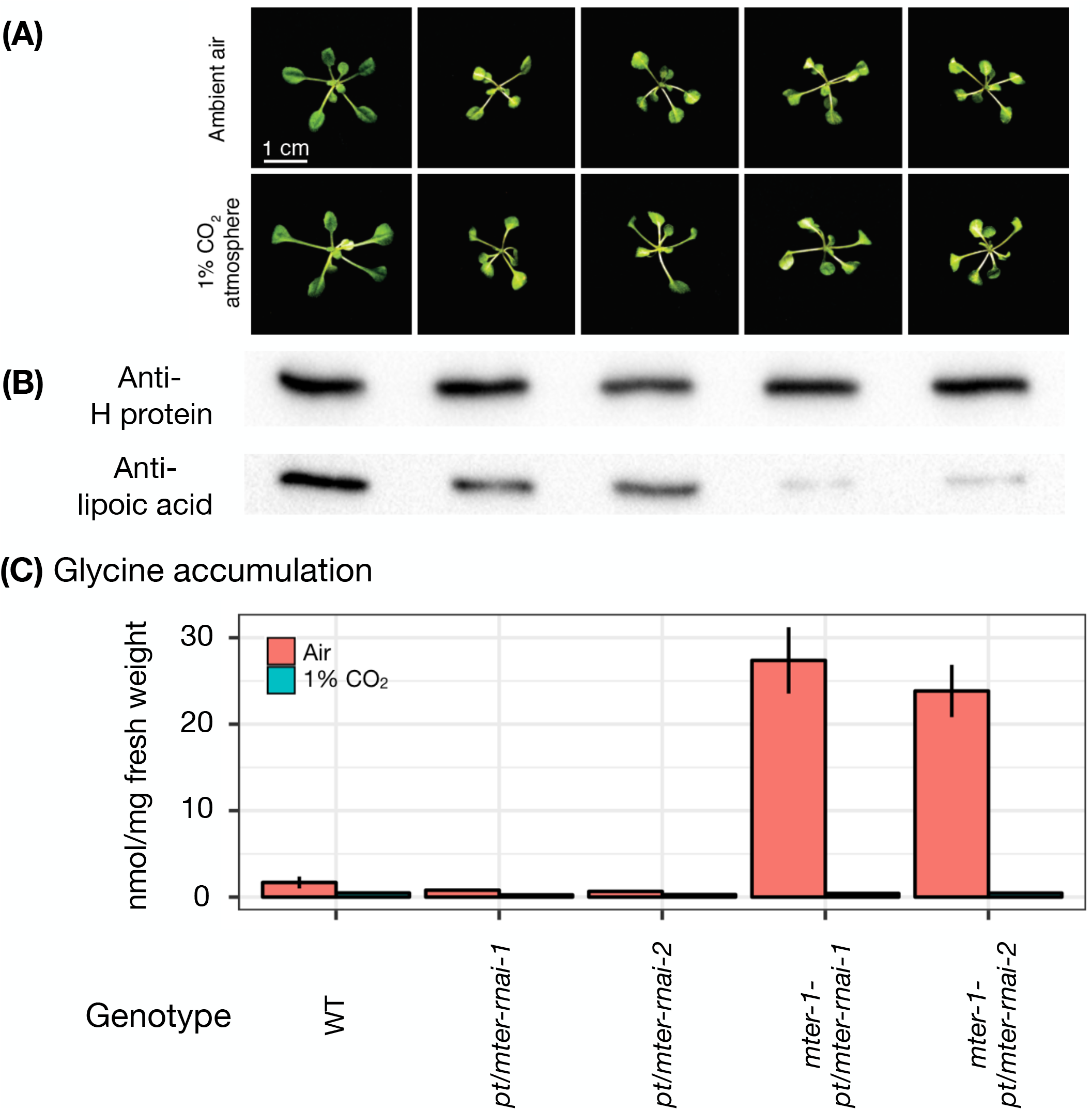
Physiological characterization of the *pt/mtER* and *mtER* genes **(A)** Morphological phenotypes of single or double mutants of the *pt/ mtER* and *mtER* genes. Plants were grown in either ambient air or in a 1% CO_2_ atmosphere. **(B)** Western blot analysis of the H subunit of GDC detected with anti H-protein antibodies, and the lipoylation status of the H-protein detected with anti-lipoic acid antibodies. Plants were grown in a 1% CO_2_ atmosphere. **(C)** Glycine accumulation. Plants of the indicated genotype were grown in either ambient air or in a 1% CO_2_ atmosphere.

In contrast, biochemical analyses that evaluated the metabolic status of these mutant plants indicate that both *mtER* and *pt/mtER* contribute to mtFAS. The evidence that supports this conclusion includes: a) western blot analysis that indicates the lipoylation states of GDC in *mter* and *pt/mter-rnai* single mutants is reduced to between 60% to 80% of the wild-type levels, and this protein is even further under-lipoylated, to about 10% of the wild-type level in the double mutant plants (Figure 4B); b) accompanied with the reduction in lipoylation status of the H protein, glycine accumulation is increased by about 15-fold in these double mutant lines; and c) this latter attribute is completely reversed when these double mutant plants are grown in the 1% CO_2_ atmosphere, which suppresses photorespiration (Figure 4C). Other changes in amino acid levels were detected in these double mutant (Supplemental Table 1). Collectively therefore, we conclude that the mtFAS system appears to be redundantly enabled by two enoyl-ACP reductases, a dual localized enoyl-ACP reductase (pt/mtER; AT2G05990) and a mitochondrially localized enoyl-ACP reductase (mtER; AT3G45770).

## DISCUSSION

### Dual-localized MCAT, KR and ER components of mtFAS

The plant mtFAS system is a central pathway that contributes substrates required for a series of essential metabolic processes, including photorespiration and biosynthesis of lipid A-like molecules (4). To date four enzymatic components of the mtFAS system have been experimentally characterized, mtKAS (16-18), mtHD (4), mtPPT (19) and mtMCS (3). In this study we identified the additional three components that are required to complete the mtFAS cycle. These three components (pt/mtMCAT, pt/mtKR and pt/mtER) were shown to localize to both plastids and mitochondria, referred to as “dual-targeted”, and thus they are contributing to both mtFAS and ptFAS capabilities. In addition, the mitochondrial ER reaction appears to be catalyzed by redundant enzymes, an NADPH-dependent mtER and a dual-targeted NADH-dependent pt/mtER. The pt/mtER and mtER belong two distinct families of enzymes; pt/mtER belongs to the short chain dehydrogenase/reductase (SDR) family, whereas mtER belonging to the medium chain dehydrogenase/reductase (MDR) family (30). Despite the distinct preference for different reducing cofactors (NADH versus NADPH), both pt/mtER and mtER display a similar chain length preference for the 2-enoyl-CoA substrates, being able to almost equally utilize both medium (10 carbon atoms) and long-chain (16 carbon atoms) acyl-CoA substrates. This latter finding is consistent with the ability of the plant mtFAS system to not only generate octanoic acid for lipoic acid biosynthesis, but also longer chain fatty acids that in plants are used in the assembly of lipid A-like molecules (31).

The dual localized mtFAS components (i.e., pt/mtMCAT and pt/mtKR) had previously been characterized as components of the ptFAS system (20,21,29), and these characterizations had established the essentiality of these components during embryogenesis. Thus, deducing that these are also components of the mtFAS system required a combination of reverse genetic and transgenic strategies. Specifically, we generated plants that were deficient in mitochondrial MCAT or KR functions, but normally expressed plastidial-localized MCAT or KR functions. This could be achieved by the fact that for these two proteins, the plastid-targeting information is encompassed in the mature protein sequence, whereas the mitochondrial-targeting information is encompassed in the N-terminal signal-peptide extension sequence. Thus, the transgenically expressed, N-terminal truncated MCAT or KR proteins were singly targeted to plastids, rather than to both the plastids and mitochondria. Therefore, these transgenically plastid-only retargeted MCAT or KR alleles rescued the embryo lethal phenotypes associated with the deficiency in ptFAS, but the recovered transgenic plants displayed phenotypes that are typical of plants lacking mtFAS functionality. These phenotypes include reduced H-protein lipoylation, elevated glycine accumulation and reduced growth; the latter two attributes being reversed by growing plants in a CO_2_-enriched atmosphere. Therefore, these findings establish that these two gene products also provide mitochondrial MCAT or KR functions, respectively.

The characterization of the mitochondrial ER component was more complex, because two loci can provide this functionality, a dual-localized pt/mtER (AT2G05990) and a singly targeted mtER (AT3G45770). Plants deficient in mitochondrial ER function were generated by combining mutants that affect both loci. Namely, a combination of a T-DNA tagged null alleles at AT3G45770, and RNAi-knock-down alleles of AT2G05990. These plants exhibit traits that are typical of a deficiency in mtFAS; namely, depleted lipoylation of the H-protein, hyperaccumulation of glycine and miniature aerial organs. All these traits appear when these plants are grown in ambient air, and glycine hyperaccumulation is reversed when they are grown in an elevated CO_2_ atmosphere.

### Dual-localization of proteins to plastids and mitochondria

An increasing number (>100) of proteins generated from a single gene locus are known to be dual-targeted to both mitochondria and plastids (32). What might be the evolutionary advantages that drive the formation of dual-targeting events? An obvious benefit is the apparent economy of maintaining only one gene instead of two. However, from an evolutionary point of view, concentrating two functions in a single gene is unusual, as the more common tendency is neo-specialization via gene duplication (33). In addition, there are clear disadvantages for dual targeted proteins. For example, the protein may not function optimally in both organelles, in terms of pH optima, concentration of substrates and co-factors, and protein-protein interactions. Moreover, a single crucial mutation could lead to the loss of these functions in both compartments.

Insights maybe provided by considering the functions of the proteins that are known to be dual-targeted. Dual-targeted proteins appear to be enriched in a few specific functional groups (33), such as DNA replication, tRNA biogenesis, protein translation and metabolic processes (e.g. fatty acid biosynthesis as demonstrated in the current study). This potential bias may reflect mechanistic constrains on dual-targeting and selection pressure that could favor dual-targeting over evolutionary timescales. These processes are all essential, and retain highly conserved functions in both organelles. This functional conservation may therefore lead to the retention of dual-targeting events by ensuring that proteins from these two organelles are functionally interchangeable.

Mechanistically, for dual-targeting between mitochondria and plastids, the targeted protein has to be recognized by the import machinery of both organelles (34). Evolutionarily, the protein import machineries in these organelles arose independently, are non-homologous, and therefore would normally be expected to recognize different targeting signals (35). Indeed, in the case of pt/mtMCAT, pt/mtKR and pt/mtER, these proteins appear to utilize bi-partite targeting signals, an N-terminal signal unique for one organelle and an internal signal that specifies import to the other organelle. Deletion of the N-terminal signal of pt/mtMCAT and pt/mtKR abolishes import into mitochondria and enhances import into plastids, whereas the contrary was observed for pt/mtER. Similar situations have been reported with many other such dual targeted proteins (e.g. AspRS, LysRS and ProRS (36,37), BT1 (38), and PGP1 (39)); deletion of their N-terminal targeting sequences only affects localization to one of these two organelles. In the case of the mtER isozyme, deletion of the N-terminal targeting sequence results in the cytosolic localization of the protein, indicating the presence of an additional signal that is needed for dual-localization.

Dual localization of pt/mtMCAT, pt/mtKR and pt/mtER, a trait acquired during the co-evolution of plastids and mitochondria, suggests that further investigations of protein sorting mechanisms and re-evaluation of organelle proteomes may be useful to reveal how generalizable this phenomenon is in plant cells. Moreover, the general persistence of such genetic and biochemical redundancy in plant metabolism may indicate evolutionary biological advantages to the generation of such complexity in metabolism, a characteristic that was initially surmised from the study of specialized metabolism (40).

## METHODS

### Yeast strains and genetic complementation

The yeast *S. cerevisiae* strain deficient in the *MCT1*, *OAR1* and *ETR1* genes (YBR026C; BY4741 background; and MATa) was obtained from Thermo Scientific. The yeast *MCT1* (primers M1-M2, see Supplemental Table 2 for primer sequences), *OAR1* (primers M3-M4) and *ETR1* genes (primers M5-M6) of wild-type BY4741 strain were cloned into YEp351 vector (PGK promoter-driven gene expression) (26). Arabidopsis *pt/mtMCAT* (AT2G30200, primers M7-M8), *pt/mtKR* (AT1G24360, primers M9-M10), *pt/mtER* (AT2G05990, primers M11-M12) and *mtER* (AT3G45770, primers M13-M14) genes of Arabidopsis Col-0 strain were cloned into YEp351M (4) (5’ plant targeting signals were replaced by the yeast *COQ3* mitochondrial presequence CDS). Complementation test was performed as previously described (27).

### Protein overexpression and in vitro kinetic assays

The *mt/ptER* (primers M15-M16) and *mtER* (primers M17-M18) genes were cloned into pBE522 vector (41). These constructs express recombinant proteins with a His-tag located at the N-terminus. Recombinant proteins were expressed in the *E. coli* BL21* strain (Invitrogen, Carlsbad, CA), and further purified using Probond Nickel-Chelating Resin (Invitrogen). The kinetic constants of mt/ptER and mtER were determined spectrophotometrically as previously described (42) with modification. Specifically, the enoyl-CoA substrates (i.e., *trans*-Δ^2^-10:1 and *trans*-Δ^2^-16:1) were synthesized as previously describes (4). Concentrations of three ingredients were optimized (i.e., 30 mM potassium phosphate (pH = 7.8), 2-200 μM enoyl-CoA, 300 μM NADPH or NADH, and 3 ug/mL recombinant proteins). The consumption of NADPH or NADH at 340 nm (for 0, 5, 10, 15 and 20 min at 22°C) was measured to monitor the enoyl-CoA reduction reaction. Kinetic values for mt/ptER and mtER were calculated using Prism version 5.0 (GraphPad Software).

### Plant strains and genetic transformations

The Arabidopsis genetic strains used in this study include: *mcat-1* (Ler background, GT_5_100190), *kr-1* (Col-3 background, SAIL_165_A11), *mter-1* (Col-0 background, SALK_056770) and *mter-2* (Col-0 background, SALK_085297), which were obtained from Arabidopsis Biological Resource Center (Columbus, OH; http://abrc.osu.edu).

In the GFP transgene experiments, PCR inserts were obtained using the following primers: *pt/mtMCAT_1-1179_* (primers M19-M20), *pt/mtMCAT_1-216_* (primers M19-M21), *pt/mtMCAT_205-1179_* (primers M22-M20), *pt/mtKR_1-957_* (primers M23-M24), *pt/mtKR_1-234_* (primers M23-M25), *pt/mtKR_214-957_* (primers M26-M24), *pt/mtER_1-1167_* (primers M27-M28), *pt/mtER_1-261_* (primers M27-M29), *pt/mtER_262-1167_* (primers M30-M28), *mtER_1-1122_* (primers M31-M32), *mtER_1-300_* (primers M31-M33), *mtER_301-1122_* (primers M34-M32), UGP3_*1-600*_(primers M35-M36). PCR inserts were cloned into pENTR/D-TOPO vector (Invitrogen), and subcloned into pEarleyGate103 (43) using Gateway LR Clonase II Enzyme Mix (Invitrogen).

In complementation transgene experiment, *pt/mtMCAT_1-1182_* (primers M19-M37), *pt/mtMCAT_205-1182_* (primers M22-M37), *pt/mtKR_1-960_* (primers M23-M38) and *pt/mtKR_214-960_* (primers M26-M38) were cloned into pENTR/D-TOPO vector, and subcloned into pEarleyGate100 (43). Destination vectors were used to transform the Arabidopsis heterozygous mutant plants deficient in *pt/mtMCAT* and *pt/mtKR*, respectively.

In the RNAi experiment, pt/mtER_202-380_ (primers M39-M40) was cloned into pENTR/D-TOPO, and subcloned into pB7GWIWG2(II) (44). Destination vectors were used to transform the Arabidopsis wild-type plants or *mter-1* mutant plants as previously described.

Plant seedlings were grown on Murashige and Skoog agar medium at 22 °C under continuous illumination (photosynthetic photon flux density 100 μmol m^−2^ s^−1^) as previously described (45). The atmospheric CO_2_ condition was maintained in ambient level or in the 1% CO_2_ level (in a growth chamber).

### Western blot

Total protein was extracted from 200 mg fresh aerial organs of plants at 16 DAI as described previously (46). Western blot analysis was carried out with anti-lipoic acid antibodies (EMD Millipore, Billerica, MA) or with anti-H protein antibodies (a gift from Dr. David Oliver at Iowa State University) on 50 μg total protein as described previously (18,19).

### Metabolomic analyses

Metabolites (3-6 replicates) were extracted from 50 mg fresh aerial organs of plants (grown in a completely randomized design) at 16 DAI, and analyzed using multiple analytical platforms: Agilent 7890 GC-MS system for fatty acids (47); Agilent 1200 HPLC system equipped with a fluorescence detector for amino acids (19); and Waters Xevo G2 Q-TOF MS equipped with Waters ACQUITY UPLC system for glycerolipids and chlorophylls (48). Log_2_-ratio and SE were calculated as described previously (49).

## ACKNOWLEDGEMENTS

The authors acknowledge Kouji Takano (RIKEN Center for Sustainable Resource Science, Yokohama, Kanagawa, Japan) for technical assistance in obtaining the lipidome data; Drs. Lloyd Sumner (University of Missouri, Columbia, MO) and Richard Dixon (University of North Texas, Denton, TX) for helpful discussions; and the WM Keck Metabolomics Research Laboratory and the Confocal Microscopy Facility (Iowa State University) for providing access to instrumentation in the metabolomic analyses and subcellular localization studies, respectively. This work was partially supported by the National Science Foundation (Awards IOS1139489, EEC0813570 and MCB0820823 to B.J.N.), the State of Iowa, the Japan Science and Technology Agency Strategic International Collaboration Research Program (SICORP), and RIKEN Pioneering Project Integrated Lipidology.

